# A drug cocktail of rapamycin, acarbose, and phenylbutyrate enhances resilience to features of early-stage Alzheimer’s disease in aging mice

**DOI:** 10.1101/2024.01.26.577437

**Authors:** Jackson Wezeman, Martin Darvas, Nadia Postupna, Jenna Klug, Ruby Sue Mangalindan, Addison Keely, Kathryn Nguyen, Chloe Johnson, Manuela Rosenfeld, Warren Ladiges

**Affiliations:** Department of Comparative Medicine, School of Medicine, University of Washington, Seattle, WA; Department of Laboratory Medicine and Pathology, School of Medicine, University of Washington, Seattle, WA

**Keywords:** Aging, Cognitive impairment, Adeno-associated viral vector, Alzheimer’s disease, Resilience, Drug cocktail, Rapamycin, Acarbose, Phenylbutyrate, Aging pathways

## Abstract

The process of aging is defined by the breakdown of critical maintenance pathways leading to an accumulation of damage and its associated phenotypes. Aging affects many systems and is considered the greatest risk factor for a number of diseases. Therefore, interventions aimed at establishing resilience to aging should delay or prevent the onset of age-related diseases. Recent studies have shown a three-drug cocktail consisting of rapamycin, acarbose, and phenylbutyrate delayed the onset of physical, cognitive, and biological aging phenotypes in old mice. To test the ability of this drug cocktail to impact Alzheimer’s disease (AD), an adeno-associated-viral vector model of AD was created. Mice were fed the drug cocktail 2 months prior to injection and allowed 3 months for phenotypic development. Cognitive phenotypes were evaluated through a spatial navigation learning task. To quantify neuropathology, immunohistochemistry was performed for AD proteins and pathways of aging. Results suggested the drug cocktail was able to increase resilience to cognitive impairment, inflammation, and AD protein aggregation while enhancing autophagy and synaptic integrity, preferentially in female cohorts. In conclusion, female mice were more susceptible to the development of early stage AD neuropathology and learning impairment, and more responsive to treatment with the drug cocktail in comparison to male mice. Translationally, a model of AD where females are more susceptible would have greater value as women have a greater burden and incidence of disease compared to men. These findings validate past results and provide the rationale for further investigations into enhancing resilience to early-stage AD by enhancing resilience to aging.

## Introduction

Aging is a complex and multifaceted process arising from the breakdown of many necessary pathways. The aging process has thus been implicated as the greatest risk factor for the onset and development of “age-related diseases.” While individual therapies may help to alleviate disease-specific phenotypes, a much broader and impactful focus is on aging itself. Zhang summarized it well stating, “The concept of geroscience is that since ageing is the greatest risk factor for many diseases and conditions, targeting the ageing process itself will have the greatest impact on human health” (Zhang 2023). Resilience aims to quantify biological aging and is defined as the ability to respond and recover from environmental stressors which disrupt homeostasis (Schorr 2018). Together, the geroscience concept and resilience could work to provide a platform to address aging and standardize evaluation and discovery toward age-related diseases. This study aims to examine the ability of resilience to delay the onset of Alzheimer’s disease cognitive and neuropathology phenotypes.

To enhance resilience to aging, a multi-target approach may be helpful to address the many pathways of aging. Previous studies have examined a drug cocktail consisting of rapamycin, acarbose and phenylbutyrate for its impact on resilience (Zhou 2022). Rapamycin targets the mTOR pathway and has been known for its ability to extend lifespan in mice as well as reduce the presence of heart disease, inflammation, and cellular senescence, even with some already recommending it be used for clinical treatment of Alzheimer’s Disease (AD) (Selvarani 2020). Acarbose is a regulator of glucose metabolism acting primarily in the intestine and targets alpha-glucosidase. Previous work in the field of Alzheimer’s research has found dysregulation of glucose as an important modulator in the progression of the disease (Dewanjee 2022). The Intervention Testing Program (ITP) is an NIA program aimed at investing the anti-aging effects of different drugs. ITP studies demonstrated combinatorial effects between rapamycin and acarbose, finding together they had a greater affect than the individual drug (Strong 2022). Phenylbutyrate, while being used primarily for urea cycle disorders, modulates histone deacetylation and acts as a chaperone for proteins in the endoplasmic reticulum. In a recent trial for Amyotrophic Lateral Sclerosis (ALS), phenylbutyrate was shown to reduce functional decline in human patients (Paganoni 2020). When the drugs are combined into a drug cocktail, it was found not only to increase resilience to aging more than any combination of two or single drug, but it was also able to modulate aging in the brain and in main body organs (Zhou 2022, 2023).

Many murine models of Alzheimer’s are unable to accurately simulate the onset and progression of the disease. One popular model, the 5xFAD mouse line, is a transgenic model using 5 amyloid precursor protein mutations to induce amyloid plaque formation in the brain. These mice, however, start developing pathology in the fetal stages. Not only does this remove the aging component from the brain but AD typically starts in humans around early to mid-life, far before cognitive phenotypes. To have better control over and to monitor the process of onset of disease phenotypes, an adeno-associated-viral (AAV) vector model was created expressing human amyloid beta-42 (Aβ-42) peptide, and P301L tau, a human mutated sequence of tau protein. There are several major advantages to this model. First, disease progression can be modeled in aging mice. Because the vector is able to be administered to mice of any age, the aging neuropathology of these mice can be allowed to interact with induced expression of these proteins for a much more translational model. Second, control of the onset additionally allows for the researcher to have greater control experimentally to assess how diet, environment, or other factors affect the induced changes. This means the model can be adapted to a number of experiments. Lastly, the landscape of AD research has begun to focus on how to treat early stages of the disease and how to prevent its onset. The control in creating a model capable of simulating the beginning stages of disease progression opens the door for development of therapies at these stages and strengthens translational value. This study propositions that increasing resilience to aging through a drug cocktail of rapamycin, acarbose, and phenylbutyrate will delay or prevent the onset and/or progression of AD neuropathology and cognitive impairment in an AAV vector model of AD. This study was designed to characterize cognitive impairment and body composition phenotypes as well as characterize the changes in neuropathology in the brains of mice to determine the ability of the AAV vector to simulate characteristic AD neuropathology, and the effect of the drug cocktail in modulating it.

## Materials and Methods

### Mice

This study used 80 male and 80 female 22-month-old C57BL/6J mice. Forty males were selected at random to receive a medicated diet consisting of rapamycin, acarbose, and phenylbutyrate with the remaining forty males receiving non-medicated feed. The females were similarly separated. Cohorts of twenty mice were formed from ten medicated fed mice and ten non-medicated fed mice, separated by sex. After 2 months of treatment, mice were infected with AAV-AD or AAV-SHAM vector, to which the primary researcher was blinded. After an additional 5 months, when mice were 27 months of age, end-of-study assays were performed followed by carbon dioxide (CO_2_) asphyxiation. Brains were immediately harvested and sectioned along the midsagittal plane into left and right halves, which were then immersed in 10% neutral buffered formalin (NBF) and allowed to fix for 72 hours before being transferred to 70% ethanol. The left sagittal hemispheres of the brain were then submitted to Alzheimer’s Disease Research Core – Neuropathology Lab at Harborview Hospital in Seattle, Washington where they were processed for paraffin embedding, serially sectioned at 5 μm into coronal slices. The main body organs were trimmed such that sections were both fixed and flash frozen. The right sagittal hemisphere was flash frozen.

### Diet

The medicated repelleted rodent chow diet contained 14 ppm encapsulated rapamycin, 1000 ppm acarbose, and 1000 ppm phenylbutyrate. The non-medicated diet consisted of only repelleted regular chow. Each drug was delivered to Test Diet Inc. (Indiana, USA) where both diets were produced. Upon arrival the diets were refrigerated and each week the diet was replaced in the cage to ensure quality. Food intake was monitored to determine the amount of food ingested. Intake was calculated each week by comparing the initial weight of food supplied to each cage to the final weight of food leftover. The consumption for each mouse was averaged across the cage. Mice were fed ad libitum.

### Vector

An AAV vector delivery system was used to induce expression of Aβ-42 and P301L tau in mouse neurons. This AAV vector is a non-replicating DNA parvovirus that has a specially designed capsid known as AAV PHP.eB, which is a subvariant of AAV9, engineered specifically to have an affinity for neurons. This variant crosses the blood brain barrier through the receptor LY6A. Subvariant AAV PHP.eB targets a specific isoform of the LY6A receptor present in significant numbers only in C57BL/6 mice. Other mouse strains do not let the virus into the brain Mathiesen 2020). There are three different constructs in total (Table 1). One induces expression of human Aβ-42, one induces P301L, a human mutated tau known to instigate AD tau pathology, and one induces Cherry, a nonsense protein. The Aβ-42 vector uses the EF-1a promoter, contains a BRI sequence to anchor the protein in the cell membrane, and contains a GFP tag for reporting. The tau vector uses the CAG promoter. The cherry vector uses the hSyn promoter.

**Table 1.** The contents of each construct including cDNA sequence, promoter, additional sequences, and reporters. Only the Aβ-42 construct has a reporter.

| cDNA Sequence | Promoter | Additional Sequence | Reporter |
| --- | --- | --- | --- |
| Human Aβ-42 | EF-1a | BRI membrane anchor | Green Fluorescent Protein (GFP) |
| P301L-Tau (FTD) | CAG | N/A | N/A |
| Cherry (Nonsense) | hSyn | N/A | N/A |

The concentration of the injected AD vector was 5 × 10^11^ Genome Copies (GC) for both the Aβ-42 and tau constructs while the SHAM vector concentration was 1 × 10^12^ GC for the cherry construct. The prepared vector was diluted with sterile filtered PBS such that each dose was 80 μl for each administration. Mice were under isoflurane anesthesia for 10 minutes before administration of the vector retro-orbitally and placed into a recovery cage immediately after. Retro-orbital administration was chosen for its low lethality rate and its ability to transduce the entire brain, in comparison to other methods such as trans-cranial and intranasal, where transduction is specific to the region injected or occipital bulb, respectively (Konno 2020, Mathiesen 2020, Hu 2021).

### QMR

Quantitative Magnetic Resonance Imaging (QMR) utilizes magnetic resonance capability in biological tissues to quantify fat and lean tissue. The machine is standardized using canola oil standard of known fat. Mice are placed in a specialized plastic tube and inserted into the machine. One scan takes 50 seconds. Three scans are completed and averaged to ensure accuracy. QMR was performed once prior to infection with the vector and once again at endpoint.

### Box Maze

Box Maze is a spatial navigation learning task developed to test cognitive function in mice (Darvas 2020). The maze consists of a rigid plastic box with 7 dud escape holes and 1 true escape hole leading to a dark empty cage. The box is lined with a reflective lining and a light is shined over the box to create a mild stressor. To start the evaluation mice are placed in the center of the maze and allowed 120 seconds to solve the maze. If the time is exceeded, mice are guided toward the exit. Once the mice are in the escape cage and the 120 seconds of the trial have passed, the mouse is returned to the home cage for 2 minutes and the process is repeated for 4 trials. It is expected mice with cognitive impairment will have longer average escape times.

### Immunohistochemistry

Brains were randomly selected high performing and low performing box maze mice representing each of the four sex/diet cohorts. This totaled 14 male and female non-medicated mice and 15 male and female medicated mice to be stained for neuropathology. Brains were stained on 4-5 μm sections with an Abcam kit (HRP/DAB Rabbit Kit: ab64261, HRP/DAB Mouse Kit: ab64259). Slides were first hydrated in xylene, 100% ethanol, 95% ethanol, 70% ethanol, and water followed by antigen retrieval by citrate buffer pH 6 or Tris-EDTA pH 9. The slides were allowed to cool at room temperature. Once cool, a hydrophobic barrier was drawn, and the slides were washed in TBST for 5 minutes. Next, the slides were incubated with a peroxidase block for 10 – 30 minutes, washed three times for 5 minutes, incubated with a protein block for 10 – 30 minutes, washed three times for 5 minutes, and incubated overnight at 4°C with primary antibody diluted in TBST. Slides were washed with TBST three times for 5 minutes, incubated with secondary antibody for 10 – 30 minutes, washed three times for 5 minutes, incubated with streptavidin for 10 – 30 minutes, washed three times for 5 minutes and incubated for 30 – 150 seconds with DAB Chromogen. After three water washes for 3 minutes, the slides were dehydrated in 70% ethanol, 100% ethanol, and xylene before having slide covers mounted with hydrophobic mounting media. The primary antibodies used are as listed: 1/1500 GFP (Abcam: ab290), 1/1000 IBA1 (Abcam: ab178846), 1/500 H31L21 (Invitrogen: 700254), 1/100 HT7 (Invitrogen: MN1000B), 1/(250 or 300) Synaptophysin (Novus: NBP2-25170), 1/200 yH2AX (Proteintech: 10856-1-AP), 1/800 MCP-1 (Novus: NBP1-07035), 1/250 PSD95 (Abcam: ab18258), 1/200 ATG5 (Invitrogen: MA5-35339), 1/2000 GFAP (Invitrogen: PA1-10019). HT7 was the only anti-mouse antibody used and a mouse-on-mouse blocking agent (Invitrogen: R37621) was applied for 30 minutes post protein block washes to reduce noise from secondary antibody binding and an additional three 5-minute washes were added before primary antibody incubation. Listing time ranges and two different antigen retrievals are to account for variation needed in fine-tuning the IHC protocol to specific antibodies. Synaptophysin was the only antibody where two concentrations were used. Non-medicated mice were stained at 1/250 while medicated cohorts were stained at 1/300. This was done due to accessibility of primary antibody. Since it was expected medicated cohorts would have greater expression of synaptophysin, they were chosen for the 1/300 dilution.

### Image Processing

Digital image analysis was performed on IHC-stained slides at 200x magnification lens using the NIS-D microscope software package (Nikon Instruments Inc.). For density measurements ImageJ was used. ImageJ is an open-source image processing software package (https://imagej.net/downloads, version 1.54d). An object-based approach to quantification was used to identify positive staining based on a binary approach consisting of present/absent DAB intensity. Images first had their background subtracted before adjusting contrast and brightness values to only include positive (present) staining. Once done, measurement options allowed for the percentage of the area in the photo represented to be recorded as percentage values. For all neuropathological evaluation, only hippocampus regions of each mouse were analyzed. Density was chosen primarily to identify range of spread specifically in the context of spatial patterns of pathology.

### Statistics

For differences between cohorts, one-way ANOVA was used to assess significance. For comparison of means between two groups, Fischer’s t-test was used. For correlations, Pearson’s coefficients were determined to have strong correlations if the coefficient was less than -0.5 or greater than 0.5. All statistics, p-values, and graphs represented were generated in GraphPad Prism version 10.1.0. Outliers were determined by (1.5 x IQR) further than the third or second quartile. Significance was determined at p < 0.05.

## Results

### Mice infected with AAV-Ab42/ptau maintained body weight similar to mice infected with AAV-SHAM

Weekly weights were used to calculate body weight between baseline and endpoint. Body fat mass was measured by QMR and performed prior to injection and again at endpoint. Food consumption was monitored weekly and the last four weeks of the study, when consumption had stabilized. Female mice infected with AAV Aβ-42/pTau (AAV-AD) and treated with the drug cocktail had a significant loss in weight (Figure 2A), while drug cocktail treated male mice with AAV-AD and AAV-Sham lost weight (Figure 2B). When comparing loss of fat, all female cohorts except the medicated AAV-SHAM demonstrated a significant decrease in fat (Figure 2C). Only the male non-medicated AAV-SHAM cohort had a statistically significant loss in fat (Figure 2D). In comparison of the male and female cohorts, it does appear male mice lost more weight in contrast to female mice who lost more fat. For the consumption of food, both male and female medicated cohorts consumed more food per mouse than mice in non-medicated cohorts (Figure 2E-2F). Interestingly, the medicated female AAV-SHAM cohort consumed more food on average in comparison to the medicated AAV-AD cohort, which led to a lower average weight loss for the AAV-SHAM cohort, possibly alluding to a dose-dependent response to the drug cocktail. Overall, though there was a sex-dependent trend in the distribution of weight loss, both male and female cohorts-maintained weight between AAV-AD and AAV-SHAM cohorts of the same feed.

**Figure 1.**
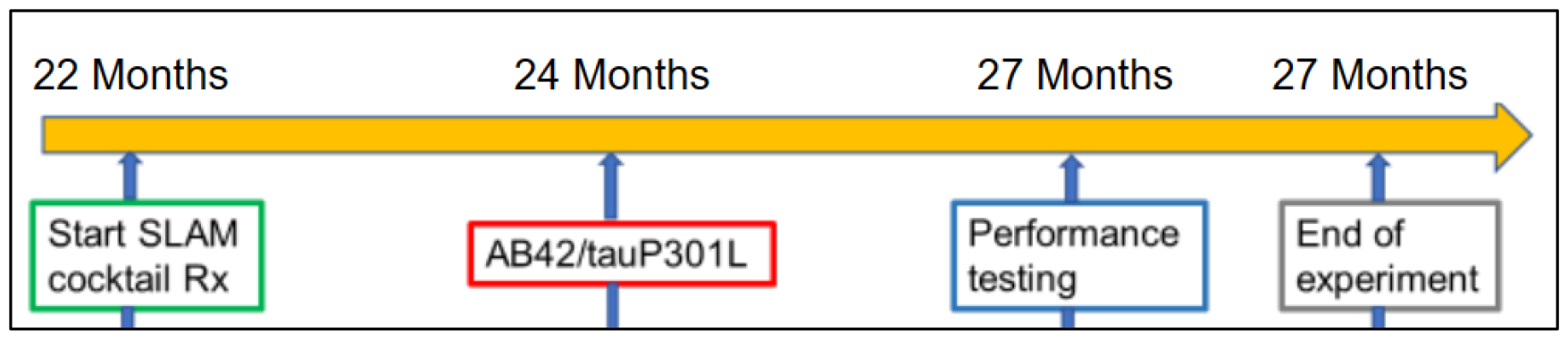
Experimental timeline. All cohorts followed this timeline and were separated by at least one week.

**Figure 2.**
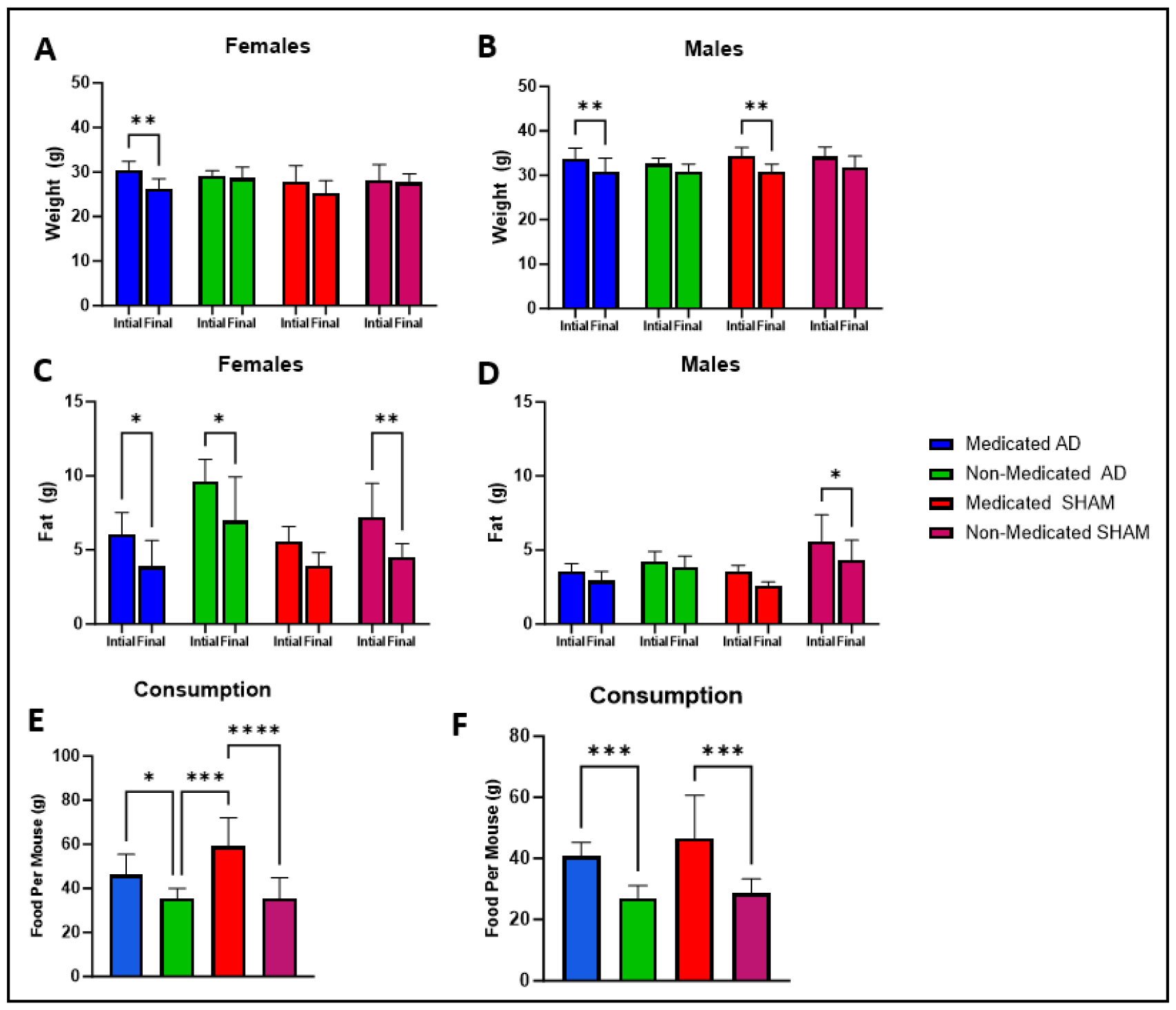
Body weight and food consumption for experimental cohorts. (A) The female medicated AAV-AD cohort lost significantly more weight compared to baseline. (B) The male medicated cohorts lost significantly more weight compared to baseline. (C) All female cohorts except medicated AAV-SHAM mice lost significant amounts of fat in comparison to pre-injection. (D) Only male non-medicated AAV-SHAM mice lost a significant amount of fat from injection to endpoint. (E) Female medicated cohorts consumed significantly more food than non-medicated cohorts. (F) Male medicated cohorts consumed significantly more food than non-medicated cohorts. n = 5-8. * = p ≤ 0.05, *** = p ≤ 0.001, **** = p ≤ 0.0001.

### The drug cocktail reduced cognitive impairment in female mice

The Box maze is a spatial navigation learning task in which mice are given 4 trials to escape the maze as quickly as possible. Each trial allowed for separation of fast and slow performers based on improvement of escape times by non-medicated AAV-SHAM mice, which served as the control. Trial 2 was selected for the female cohorts as the female control cohort continually improved their times with Trial 2 having the greatest change in escape time. Trial 3 was selected for the male cohorts as it was the last trial to mark an improvement of the control cohort on escape time. For the female cohorts, the medicated AAV-AD mice performed faster than the non-medicated AAV-AD mice regardless of injection and even when separated based on performance (Figure 3A, 3C). In male cohorts, while the non-medicated AAV-SHAM cohort had the fastest escape times, both cohorts of fast medicated mice performed faster than the fast non-medicated AAV-AD cohort (Figure 3B, 3D).

**Figure 3.**
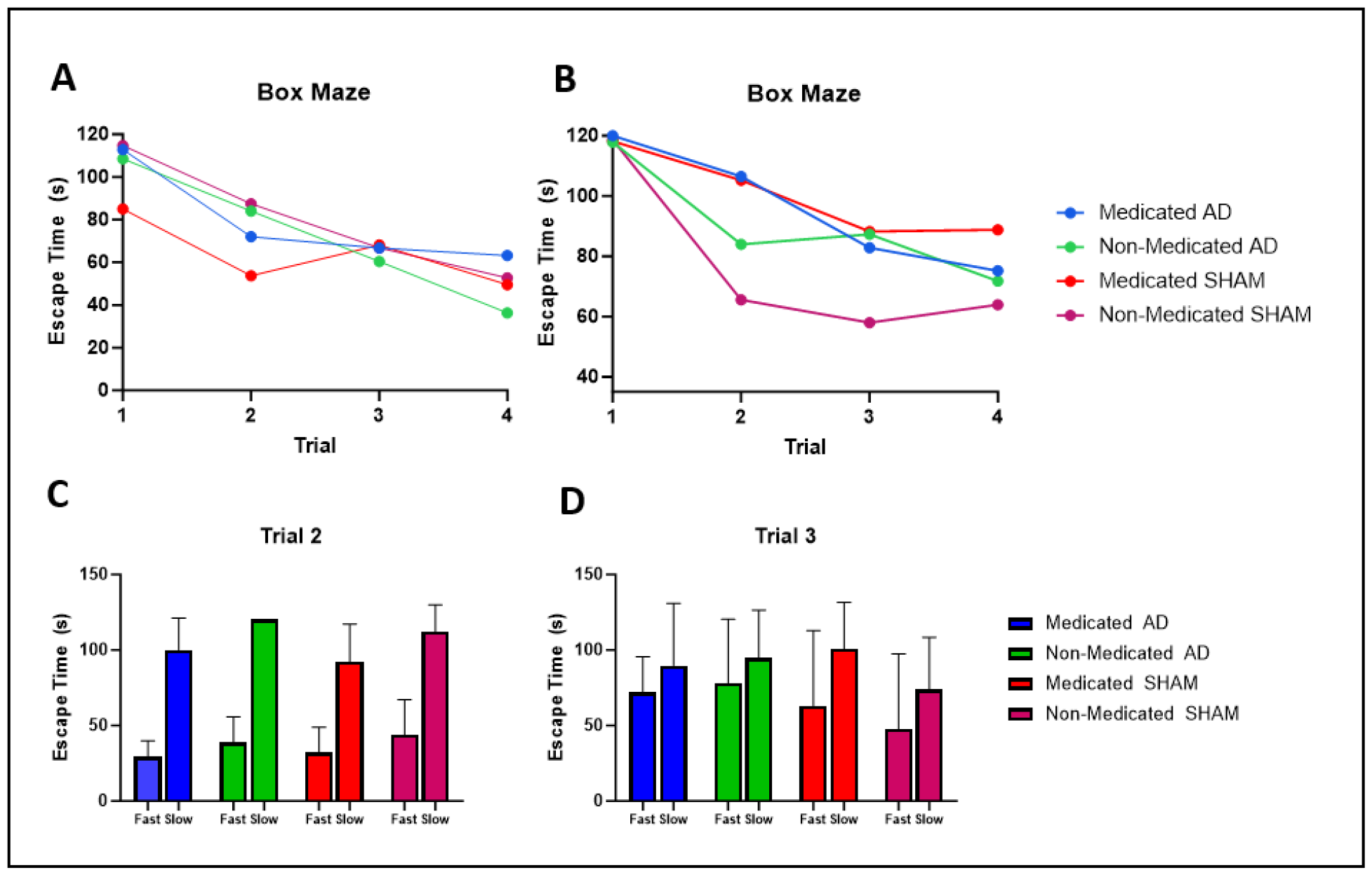
Assessment of cognitive phenotype. (A) Medicated cohorts had faster escape times than non-medicated cohorts in trial 2. (B) Male medicated cohorts performed slower than non-medicated cohorts in trial 2. (C) Separation of fast and slow performers for each female cohort for trial 2. Medicated cohorts outperform non-medicated cohorts in fast and slow separated groups. (D) Separation of fast and slow performers for each male cohort for trial 3. Non-medicated AAV-SHAM mice outperformed all other cohorts. Fast medicated cohorts performed faster than the fast non-medicated AAV-AD cohort. n = 5-8.

The Box maze data suggests a reduction in cognitive impairment for female cohorts based on small differences in fast groups and larger differences in slow groups. It was also seen that slow AAV-AD cohorts performed worse than slow AAV-SHAM cohorts. As for the male cohorts, not only did they perform slower than the female mice, but the differences in improvement were also much smaller, suggesting more severe cognitive impairment than females. It was seen that in fast groups, AAV-AD mice performed worse than AAV-SHAM mice. These results point to a rescue of cognitive impairment in female cohorts with a smaller impact on male cohorts.

### The drug cocktail suppressed expression of Aβ-42 in the hippocampus of female mice infected with the AAV-AD vector

Histopathological assessment of the brain focused on the hippocampus as the sub anatomic region of interest for its translational relevance as the primary learning and memory center adversely affected by AD in human patients. Following IHC, quantification was performed using ImageJ, an image-focused research software package, using images taken of DAB-Chromogen-stained brains. To quantify staining, density was calculated by highlighting positive pathology and measuring the “percent area” highlighted in the image. Green fluorescent protein (GFP) was used as a reporter sequence in the Aβ-42 viral construct to validate the presence of the AAV vector in neurons. IHC using an optimized anti-GFP antibody was employed. In female and male cohorts, significant differences were observed between AAV-AD and AAV-SHAM infected mice (Figure 4A, 4B). No differences in densities of GFP were seen between medicated and non-medicated AAV-AD cohorts for both sexes suggesting the drug cocktail did not interfere with viral-mediated transduction.

**Figure 4.**
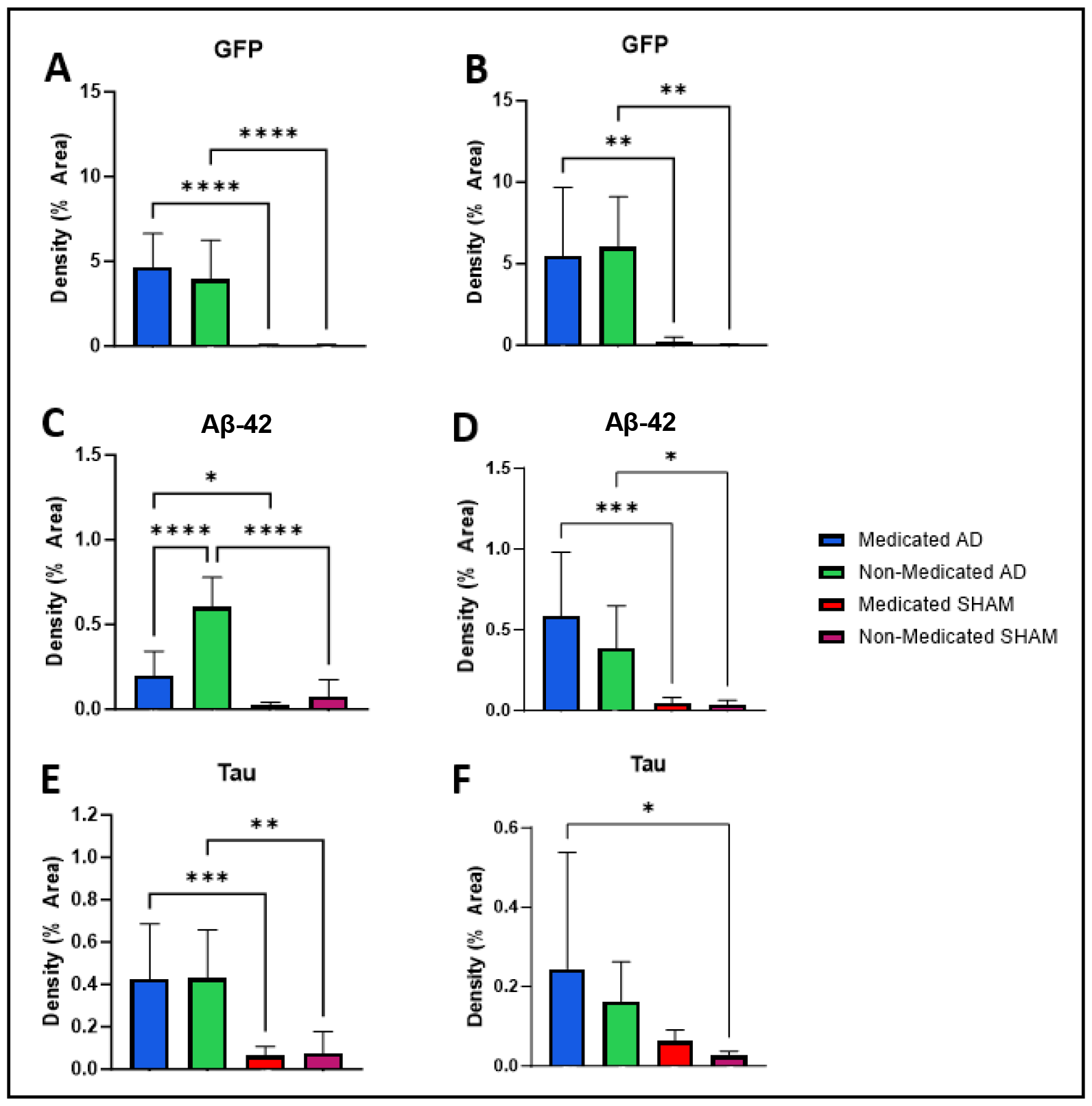
AAV-AD induced proteins GFP, Aβ-42, and tau in the hippocampus quantified in ImageJ by area density of DAB. GFP is a reporter gene for the Aβ-42 construct. Aβ-42 was stained with H31L21 which is specific for the sequence. Tau was stained by HT7 which is specific for human tau. (A-B) Both females and males had no difference between AAV-AD cohorts but significant differences between AAV-AD and AAV-SHAM infected cohorts. (C-D) Equivalent comparisons to Figure A and B, show similar trends with AAV-AD cohorts expressing significantly more Aβ-42 than AAV-SHAM cohorts. Female medicated AAV-AD mice did show a significant decrease in the density of Aβ-42 when compared to the non-medicated AAV-AD cohort. (E-F) Female cohorts in similar comparisons to Figure A and B saw similar results with AAV-AD cohorts expressing more tau than AAV-SHAM cohorts. Male cohorts saw higher averages for AAV-AD cohorts but only the medicated cohort was significant against the non-medicated AAV-SHAM cohort. n = 5-8. * = p ≤ 0.05, ** = p ≤ 0.01, *** = p ≤ 0.001, **** = p ≤ 0.0001.

As expected, expression of Aβ-42 reflected trends seen in densities of GFP. For both males and females, AAV-AD cohorts had significantly greater expression of Aβ-42 in comparison to AAV-SHAM cohorts. (Figure 4C-4D). More importantly, the female medicated AAV-AD mice had significantly less Aβ-42 expression compared to the non-medicated AAV-AD. Female AAV-AD cohorts also had significantly greater expression of tau in comparison to AAV-SHAM cohorts. No decrease in expression of tau was observed between AAV-AD cohorts (Figure 4E). While not significant against female cohorts, male expression of tau for AAV-AD infected mice was lower than female AAV-AD cohorts. While averaging higher, only medicated AAV-AD mice were significant against AAV-SHAM non-medicated mice (Figure 4F). Many males infected with the AAV-AD vector had density levels similar to AAV-SHAM infected mice. It is possible male mice may not be as susceptible as female mice to the accumulation of AD proteins as male mice appear to have less expression overall, with similar expression of GFP.

Figure 6 shows characteristic structures of AAV-AD expressed proteins in the hippocampus. Figures 5A-5C are of GFP expression, Figures 5D-5F are of Aβ-42, and Figures 5G-5I show tau expression. For each set of three images: first is an AAV-SHAM infected mouse, second is AAV-AD infected mouse, and third is a close-up of an AAV-AD infected neuron showing positive staining. In all AAV-SHAM images, no density is observed. In all AAV-AD images, examples of positive stain are circled in red. Of those circles, one has been selected for the close-up. Each enlarged figure demonstrates intercellular expression of relevant proteins and the spreading of those proteins down the axon.

**Figure 5.**
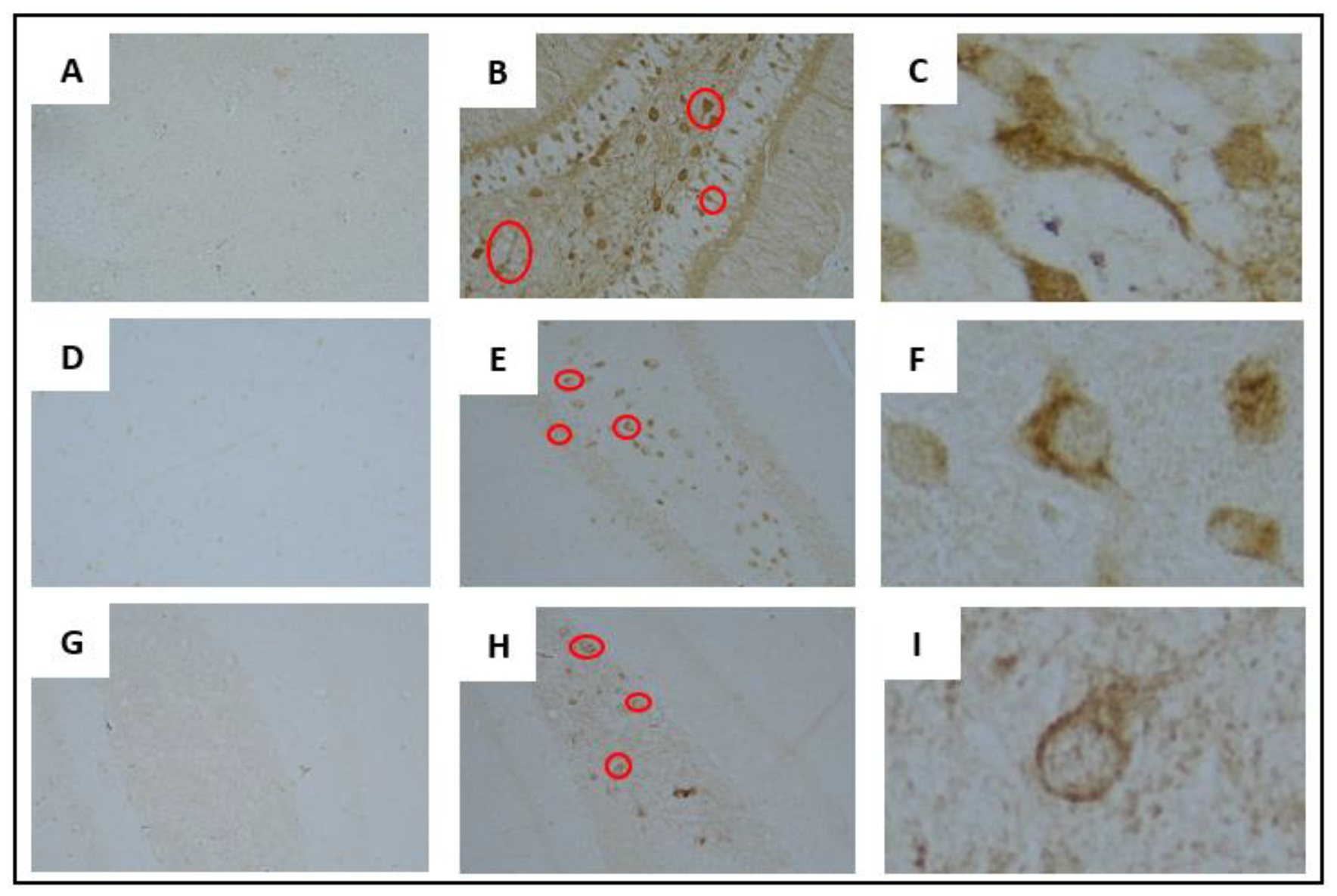
Images of immunohistochemistry stains to visualize AAV-AD protein expression in the hippocampus taken at 20x. Positive stains have annotations highlighting examples of positive staining. (A-B) Characteristic density of GFP in AAV-SHAM and AAV-AD infected mice, respectively. (C) Enlarged image of GFP positive neuron. (D-E) Characteristic density of Aβ-42 in AAV-SHAM and AAV-AD infected mice, respectively. (F) Enlarged image of Aβ-42 positive neuron. (G-H) Characteristic density of human tau in AAV-SHAM and AAV-AD infected mice respectively. (I) Enlarged image of human tau positive neuron.

**Figure 6.**
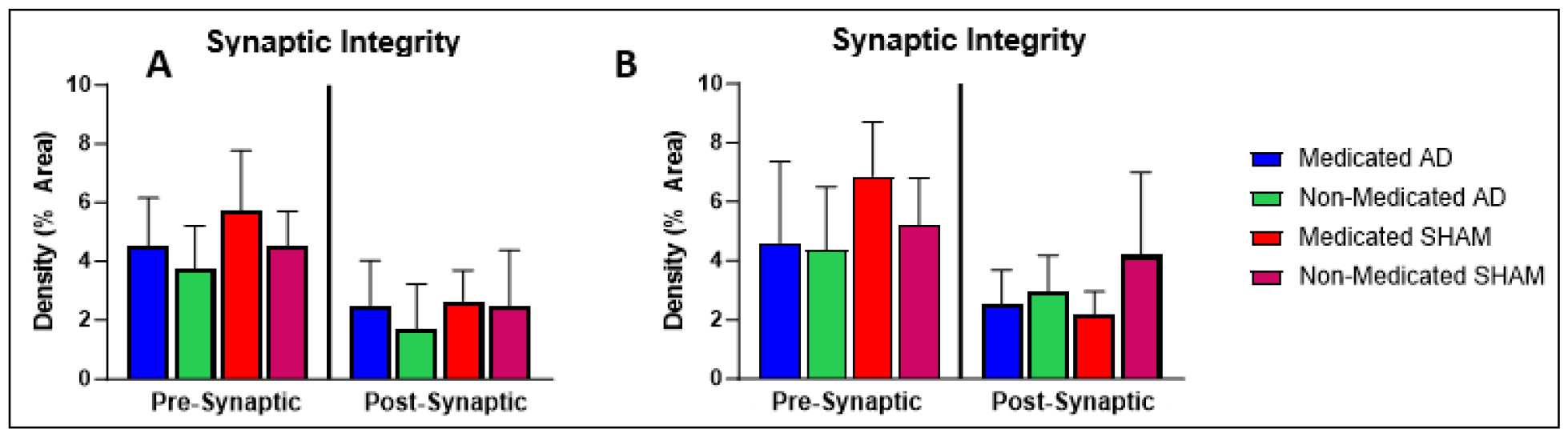
Density of characteristic markers of synaptic health in the hippocampus quantified using ImageJ. (A) Female cohorts saw higher density of synaptophysin in medicated cohorts and a decrease in synaptophysin in AAV-AD infected cohorts. A slight increase in density was observed for PSD-95 in medicated mice. (B) Male cohort saw an increase in density of synaptophysin in medicated cohorts and an overall decrease in AAV-AD infected cohorts. No differences were observed in PSD-95. n = 5-8.

### The drug cocktail altered aspects of neuropathology and concentration of non-neuronal cells in the hippocampus of mice

Because synaptic integrity is one of the first processes damaged in AD resulting in cognitive impairment, synaptic markers were analyzed in the hippocampus. Two targets were chosen, one for presynaptic integrity and one for postsynaptic integrity evaluation. Synaptophysin was chosen for its role in pre-synaptic vesicle release. PSD-95 was chosen for its role in regulation and organization of synapses. Female medicated cohorts had greater expression of synaptophysin in comparison with non-medicated cohorts with the same vector (Figure 6A). In addition, female AAV-AD cohorts had on average less synaptophysin expression in comparison to AAV-SHAM cohorts. While synaptophysin expression for the male medicated AAV-AD cohort was higher than the non-medicated AAV-AD cohort, greater differences were seen between the medicated AAV-SHAM cohort and non-medicated AAV-SHAM cohort (Figure 6B). In post-synaptic marker PSD-95 female cohorts had a slightly greater expression in medicated cohorts, while male medicated cohorts had slightly less expression than non-medicated cohorts (Figure 6A-6B). This data suggests the drug cocktail enhances synaptic integrity in female cohorts and pre-synaptic integrity in male cohorts. In addition, it also suggests mice infected with the AAV-AD vector have a reduction in synaptic integrity in comparison to their AAV-SHAM counterparts.

Figure 8 shows IHC images of staining done for pre-synaptic and post-synaptic proteins. Figure 7A demonstrates positive staining for synaptophysin with Figure 7B showing an enlarged image. Synaptophysin is expressed in the axon and appears clustered and follicular due to its direct involvement with vesicle membranes. Figure 7C demonstrates positive staining for PSD-95 with Figure 7D showing an enlarged image. PSD-95 is expressed closer to the cell body where excitatory neurons are located. PSD-95 acts as a structural mediator for bringing membrane proteins and relevant signals together. It also appears follicular, as demonstrated by small cavities of DAB staining.

**Figure 7.**
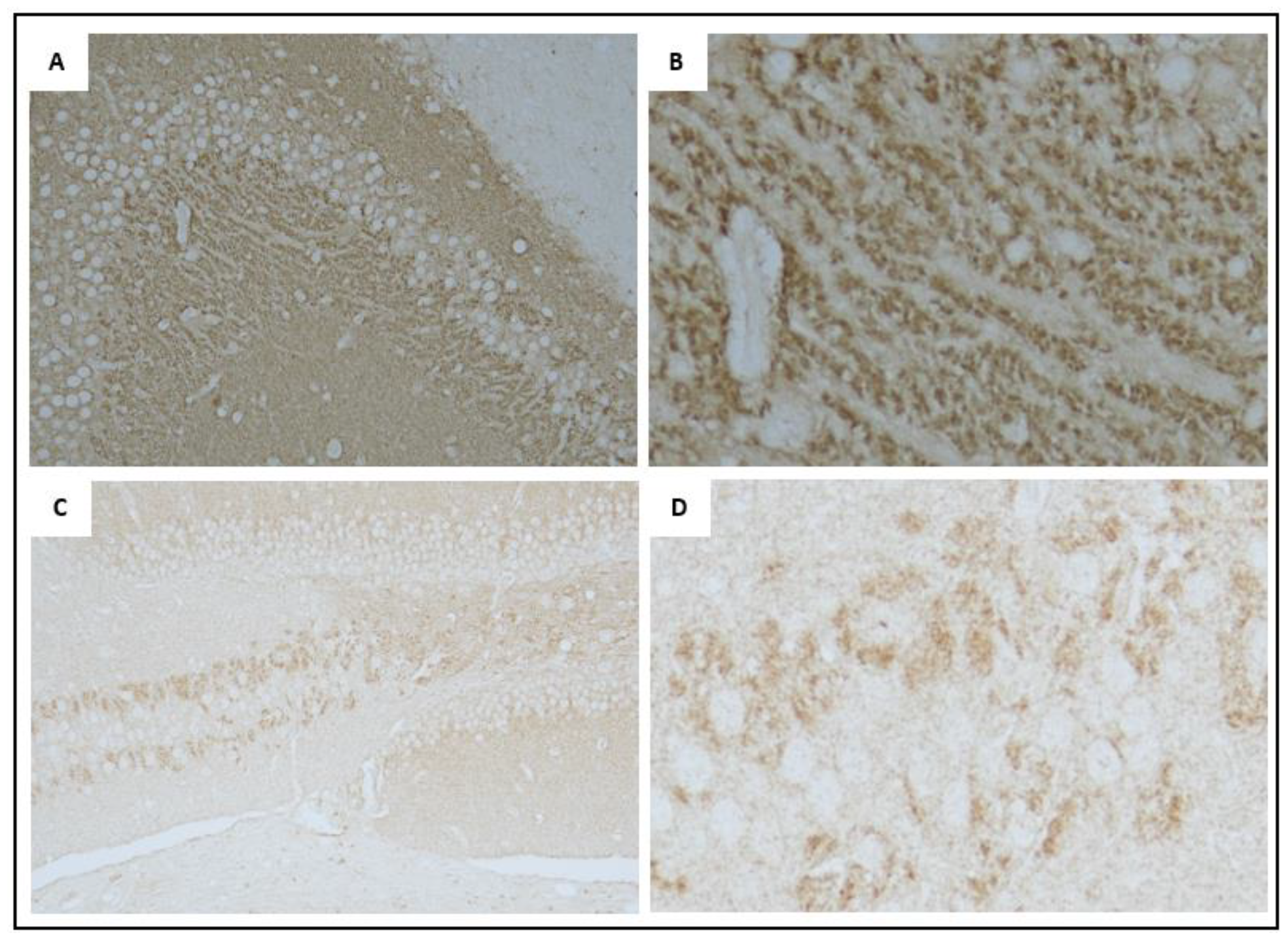
Images are representative of the characteristic markers of synaptic integrity in the hippocampus. (A) 20x image of the hippocampus. Stained for Synaptophysin. (B) Enhanced image of 8A (C) 20x image of the hippocampus. Stained for PSD-95. (D) Enhanced image of 8C.

**Figure 8.**
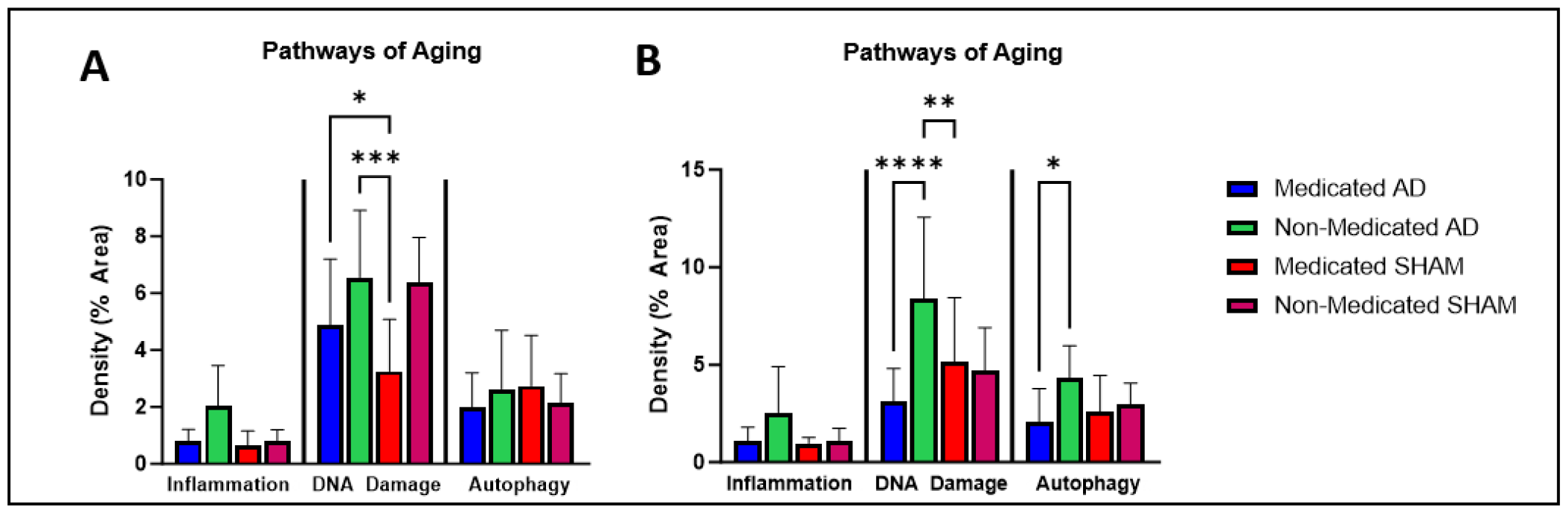
Density of IHC staining for characteristic markers of pathways of aging in the hippocampus. (A) Female AAV-AD infected mice had higher densities of MCP-1 in comparison to AAV-SHAM infected mice, as well as medicated cohorts having lower densities than their counterparts. Female AAV-AD infected mice had increased density of yH2AX compared to medicated AAV-SHAM infected mice. On average, medicated cohorts had lower densities than their non-medicated counterparts. No trends were observed for ATG5 in female mice. (B) Male AAV-AD infected mice had higher densities of MCP-1 in comparison to AAV-SHAM infected mice, as well as medicated cohorts having lower densities than their counterparts. Male non-medicated AAV-AD infected mice had increased density of yH2AX compared to medicated AAV-SHAM and AAV-AD infected mice. On average, medicated cohorts had lower density compared to non-medicated cohorts. Male medicated cohorts on average had lower density of ATG5 in comparison to non-medicated cohorts, with a significance decrease between AAV-AD infected cohorts. n = 5-8. * = p ≤ 0.05, ** = p ≤ 0.01, *** = p ≤ 0.001, **** = p ≤ 0.0001. Aging pathway biomarkers: MCP-1 for inflammation; γH2Ax for DNA damage response; ATG5 for autophagy.

These results suggest the drug cocktail influences presynaptic integrity in both males and females while in AAV-AD infected mice, expression of synaptophysin is decreased. Post-synaptic integrity seems to be affected in AAV-AD cohorts for female mice but is less affected in male medicated cohorts. The sex differences in post-synaptic integrity appear to mirror expression of AAV-AD induced protein expression. This suggests, post-synaptic integrity may be more closely linked to AD neuropathology than pre-synaptic integrity in the early stages.

As Aβ-42 and Tau expression progresses, accumulation of excess unwanted protein can lead to signals for inflammation, increases in the amount of DNA damage, and an increase in autophagy (Currais 2017, Ainslie 2021). These pathways were analyzed by IHC in the hippocampus. MCP-1 is a signal for inflammation and acts as a recruiter of additional inflammatory factors. yH2AX is a signal for DNA damage response and is upregulated in the presence of double-stranded DNA breaks. ATG5 is a marker for autophagy and is important for destruction of damaged cellular components.

In general, female AAV-AD mice, with or without drug cocktail treatment, had higher expression of representative markers in comparison to their AAV-SHAM cohorts. In addition, medicated cohorts had lower expression than non-medicated cohorts though not significant (Figure 8A). In male cohorts the same trends with AAV-AD medicated and AAV-AD non-medicated mice were observed against their counterparts, as well as the same trend between medicated and non-medicated cohorts (Figure 8B). More specifically, both medicated and non-medicated female AAV-AD cohorts had significantly higher expression of yH2AX in comparison to the non-medicated AAV-SHAM cohort. In addition, compared to their non-medicated counter parts, medicated cohorts had on average a lower density of yH2AX (Figure 8A). Male cohorts mirrored significantly less density of yH2AX in the medicated AAV-AD compared to the non-medicated AAV-AD cohort. Non-medicated AAV-AD mice also had significantly denser yH2AX than medicated AAV-SHAM mice (Figure 8B). Males did not have the same cocktail treatment trends, as only the medicated AAV-AD cohort had less density than its non-medicated AAV-AD counterpart. No significant differences in female cohorts were observed in evaluation of autophagy. Male cohorts saw a significant decrease in expression between the medicated AAV-AD compared to the non-medicated AAV-AD cohort. While not significant, the medicated AAV-SHAM also had less dense expression compared to the non-medicated AAV-SHAM cohort (Figure 8B).

Figure 9 shows images of IHC staining for inflammation, DNA damage response, and autophagy. Figure 9A demonstrates positive staining for MCP-1 with Figure 9B showing an enlarged image. MCP-1 is made in the cytoplasm and released extracellularly as a chemokine, thus evidence of staining inside and outside neurons.

**Figure 9.**
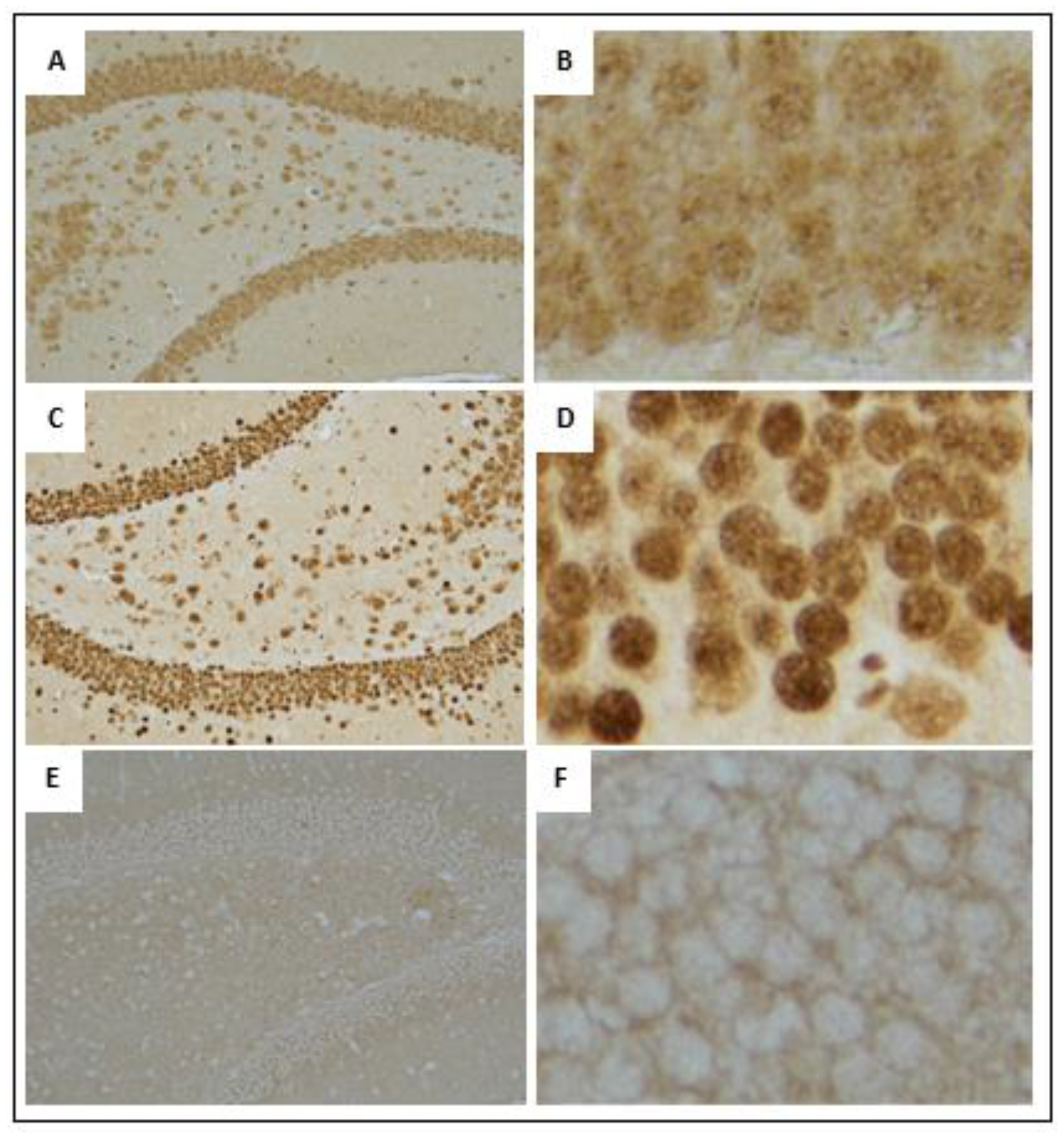
Images are representative of the characteristic markers of aging pathways in the hippocampus. (A) 20x image of the hippocampus. Stained for MCP-1. (B) Enhanced image of 10A. (C) 20x image of the hippocampus. Stained for yH2AX. (D) Enhanced image of 10C (E) 20x image of the hippocampus. Stained for ATG5. (F) Enhanced image of 10E.

Figure 9C demonstrates positive staining for yH2AX which acts on histone targets recruiting additional proteins to initiate a DNA damage response. It is localized in the nucleus and recognized as a “polka-dot” pattern. Figure 9D shows an enlarged image. Figure 10E demonstrates positive staining for ATG5 with Figure 10F showing an enlarged image. ATG5 can be localized to the nucleus but is primarily expressed in the cytoplasm where it forms complexes necessary for phagosomes. Thus, locational expression can be apparent in the axons of the neuron and outside the nucleus.

**Figure 10.**
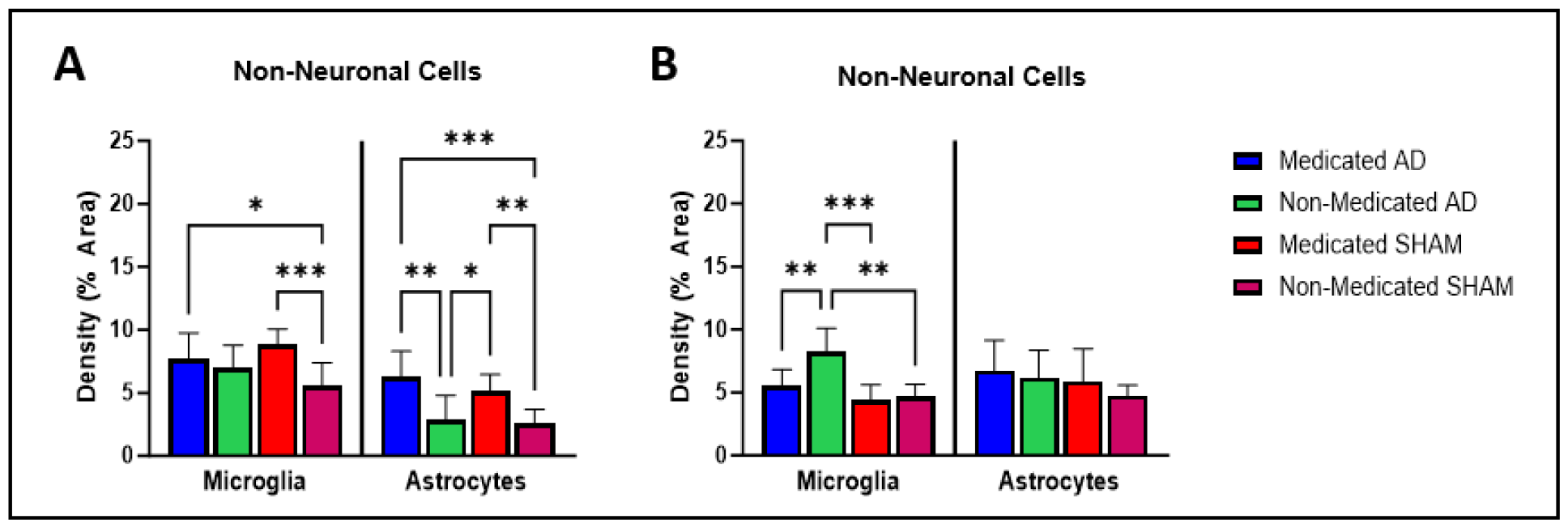
Density of IHC staining for non-neuronal cells in the hippocampus. (A) Female medicated AAV-AD and AAV-SHAM cohorts had significantly higher IBA1 density than the non-medicated AAV-SHAM cohort and higher densities compared to the non-medicated AAV-AD cohort. Both medicated AAV-AD and medicated AAV-SHAM cohorts had high densities of GFAP in comparison to both the non-medicated AAV-SHAM and non-medicated AAV-AD cohorts. (B) The male non-medicated AAV-AD cohort had significantly denser IBA1 than every other male cohort. Male AAV-AD cohorts had higher densities of GFAP in comparison to AAV-SHAM cohorts. n = 5-8. * = p ≤ 0.05, ** = p ≤ 0.01, *** = p ≤ 0.001, **** = p ≤ 0.0001.

The expression of these markers demonstrates cocktail mediated responses to inflammation and DNA damage. For both sexes, differences in inflammation between cohorts mirrored differences in DNA damage and for males, differences in autophagy. This suggests specific responses to expression of Aβ-42 and Tau, which are alleviated by the drug cocktail.

The role of non-neuronal cells in the progression of AD is significant. Microglia responses can become increased to the point of synapse engulfment and development of tau pathology (Hansen 2018). Astrocytes help in neuronal maintenance of synapses and clearing of debris but have been implicated in being activated by microglia and contributing to inflammation during AD (Orre 2014). IBA1 is a marker for activated microglia and GFAP is a marker for activated astrocytes. Surprisingly, both female medicated AAV-AD and medicated AAV-SHAM cohorts had significant increases in IBA1 expression in comparison to the non-medicated AAV-SHAM cohort. Both cohorts also had increased expression compared to the medicated AAV-AD cohort (Figure 10A). In direct contrast, for male cohorts there were no differences in IBA1 staining density except that all cohorts had significantly less IBA1 expression in comparison to the non-medicated AAV-AD cohort. It appeared on average that AAV-AD cohorts had higher expression of IBA1 than AAV-SHAM cohorts (Figure 10B). In both medicated AAV-AD and AAV-SHAM female cohorts there was a significant increase in GFAP expression than in both non-medicated cohorts (Figure 10A). Male cohorts did not follow this trend. In males, AAV-AD cohorts on average had higher expression of GFAP in comparison to AAV-SHAM cohorts. Microglia activate astrocytes and sex differences have been observed in astrocyte activation (Crespo-Castrillo 2020, Posillico 2021), which may explain the differences in non-medicated cohorts. This suggests the drug cocktail impacted astrocyte activation preferentially in female cohorts, and potentially given the intensity of the male response, in male cohorts.

Figure 11 shows images of IHC staining for non-neuronal cells. Figure 11A demonstrates positive staining for IBA1 with Figure 11B showing an enlarged image. Figure 11C demonstrates positive staining for GFAP with Figure 11D showing an enlarged image. Both cell types can be identified by the dark globular nucleus with thin wiry arms stretching out. Differences in the presence of microglia and astrocytes suggest the AAV-AD vector enhances the activation of non-neuronal cells in female mice. However, the male medicated cohorts appeared to have less microglia and more astrocytes when compared to non-medicated cohorts with the same vector, supporting a drug cocktail effect on activated astrocytes.

**Figure 11.**
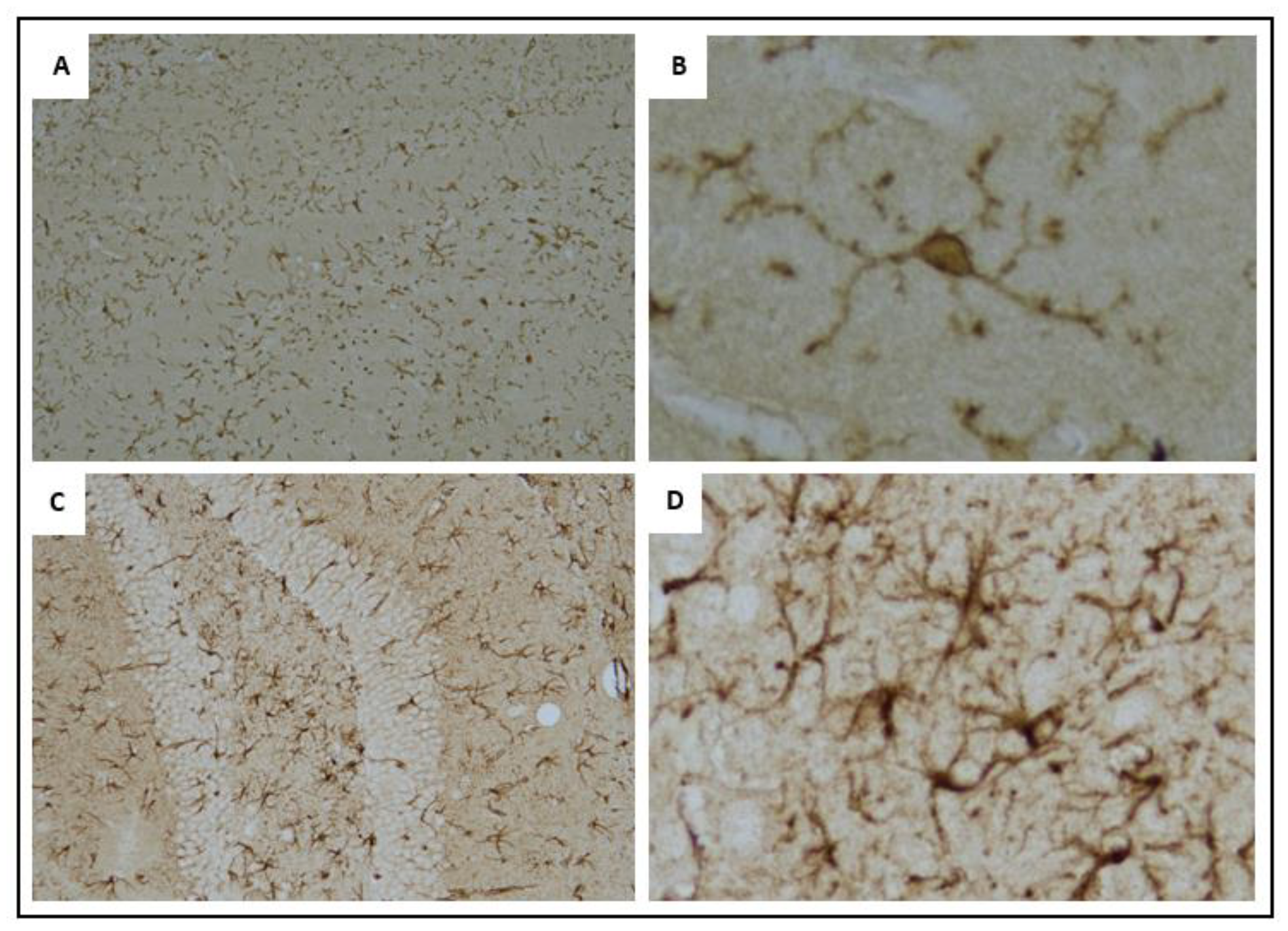
Images are representative of the characteristic markers of non-neuronal microglia and astrocytes in the hippocampus. (A) 20x image of the hippocampus, stained for microglia using IBA1 antibody. (B) Enhanced image of 12A. (C) 20x image of the hippocampus, stained for astrocytes using GFAP antibody. (D) Enhanced image of 12C.

### Associations with inflammation and autophagy appeared to be mediated by the drug cocktail in both male and female cohorts

Figures 12 and 13 contain correlation matrices comparing neuropathology, and box maze cognition data to draw connections based on variable distribution within the data sets. Associations were considered strong if the Pearson correlation coefficient was less than or equal to -0.5 and greater than or equal to +0.5.

**Figure 12.**
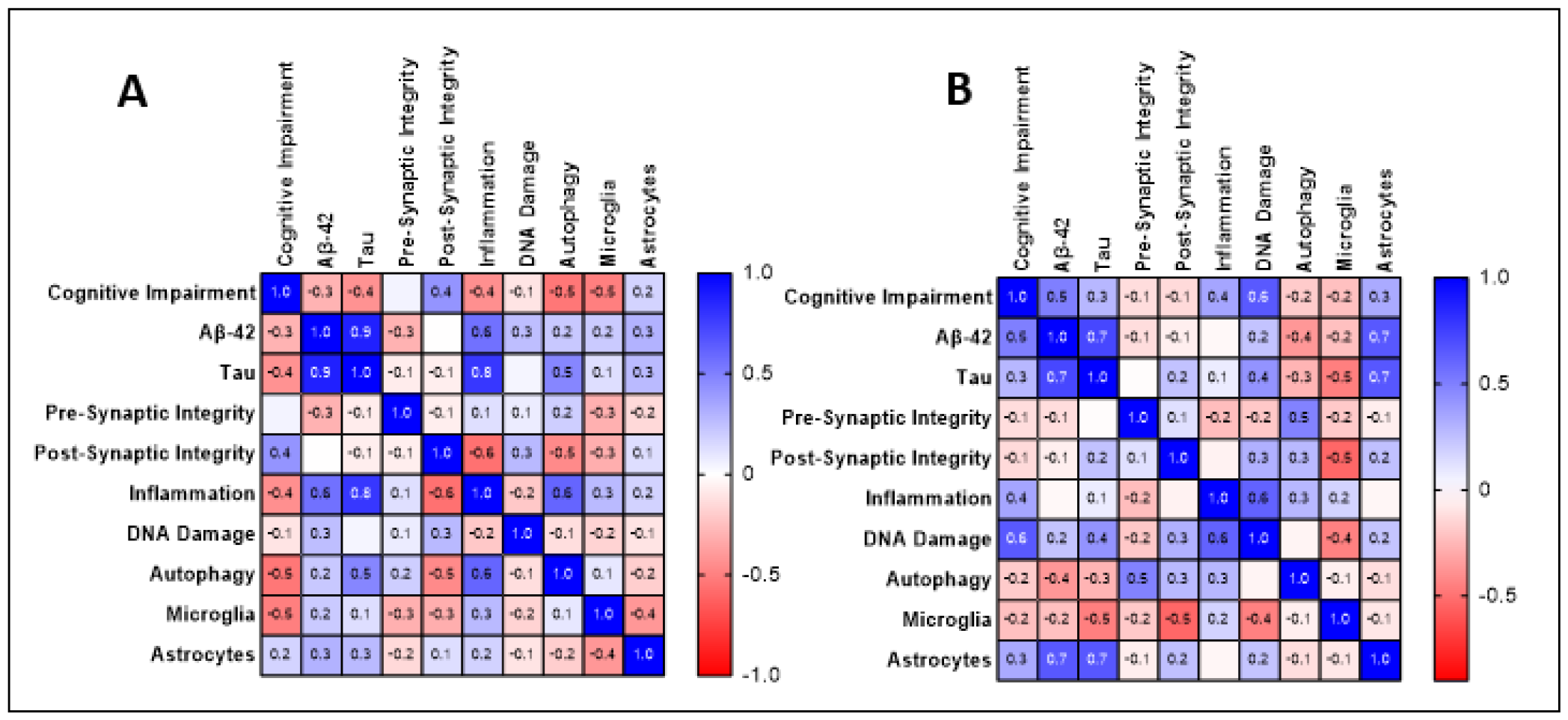
Females. Associations between neuropathology, and cognitive impairment. Numbers shown are Pearson correlation coefficients. (A) Female non-medicated cohorts show the strongest influence from inflammation, autophagy, and tau. (B) Female medicated cohorts show strong influence from Aβ-42, tau, and microglia. Strong trend set at coefficient ≥ |0.5|. n = 14-15.

**Figure 13.**
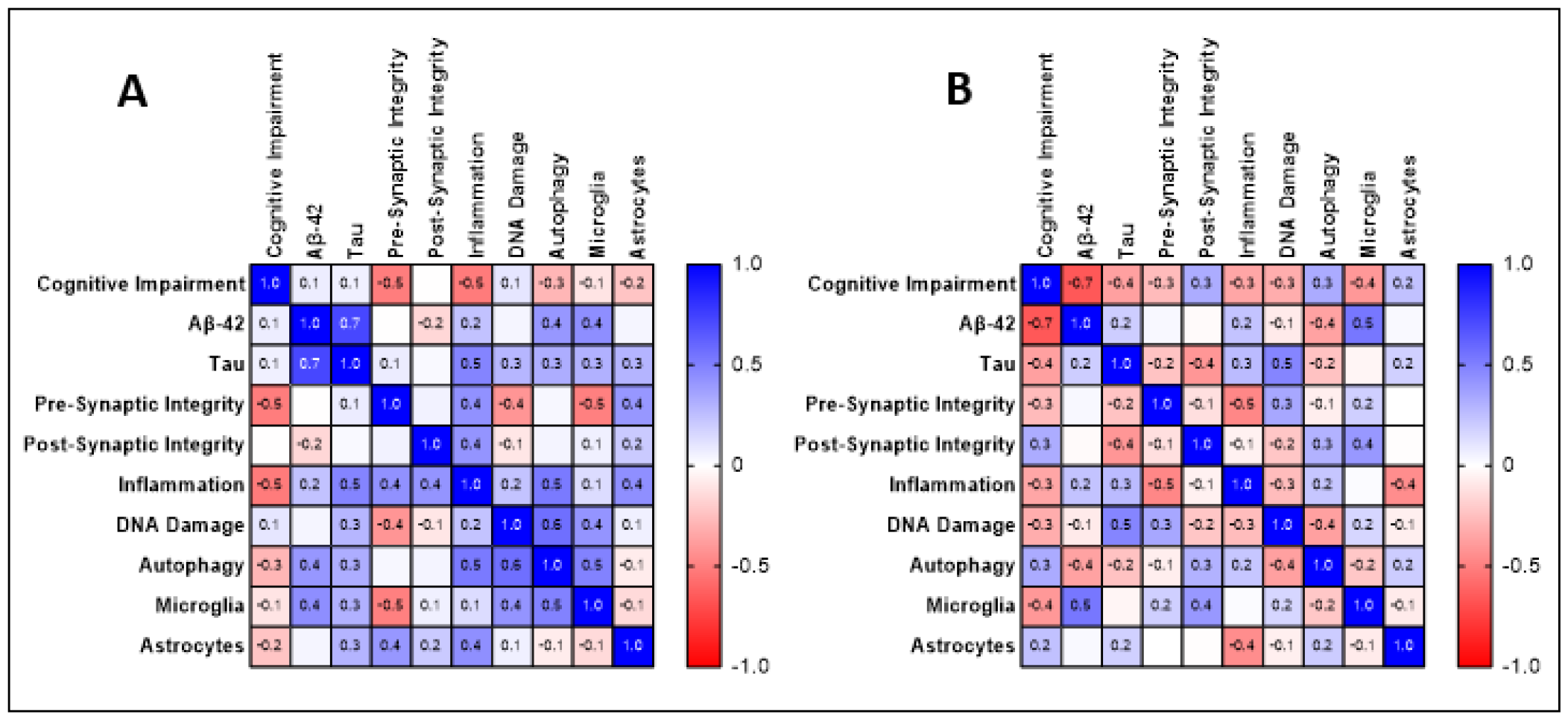
Males. Associations between neuropathology, and cognitive impairment. Numbers shown are Pearson correlation coefficients. (A) Male non-medicated cohorts show the strongest influence from inflammation, autophagy, and microglia. (B) Male medicated cohorts show strong influence from Aβ-42, and microglia. Strong trend set at coefficient ≥ |0.5|. n = 14-15.

For female non-medicated cohorts (Figure 12A), a negative strong association was seen between cognitive impairment and both autophagy and microglia. Aβ-42 density had strong positive associations with tau and inflammation. Tau had strong positive associations with inflammation and autophagy. Post synaptic integrity had strong negative associations with inflammation and autophagy. Lastly, inflammation had a strong positive association with autophagy. For female medicated cohorts in Figure 12B, cognitive impairment had a strong positive association with Aβ-42 and DNA damage. Aβ-42 had strong positive associations with tau and astrocytes. Tau had strong positive associations with astrocytes and strong negative associations with microglia. Presynaptic integrity had strong positive associations with autophagy. Postsynaptic integrity had a strong negative association with microglia. Lastly, inflammation had a strong positive association with DNA damage.

In the non-medicated cohorts (Figure 12A), the most influential variables seemed to be inflammation, autophagy, and tau. In the medicated cohorts (Figure 12B) the most influential variables seemed to be Aβ-42, tau, and microglia. This suggests the drug cocktail has relieved the burden of inflammation and autophagy, while increasing the activation of astrocytes in relation to Aβ-42 and tau, which seem to have a stronger influence over cognitive impairment.

For male non-medicated cohorts (Figure 13A), box maze had strong negative associations with presynaptic integrity and inflammation. Aβ-42 had a strong positive association with tau. Tau had a strong positive association with inflammation. Pre-synaptic integrity had a strong negative association with microglia. Inflammation had a strong positive association with autophagy. DNA damage had a strong positive association with autophagy. Lastly, autophagy had a strong positive association with microglia. For male medicated cohorts (Figure 13B), cognitive impairment had a strong negative association with Aβ-42. Aβ-42 had a strong association with microglia. Tau had a strong association with DNA damage. Lastly, presynaptic integrity had a strong negative association with inflammation.

In the non-medicated cohorts, the most influential variables seemed to be inflammation, autophagy, and microglia. In the medicated cohorts, the most influential variables seemed to be Aβ-42, microglia. This suggests that while the drug cocktail can reduce the burden of inflammation and autophagy in male mice, cognitive impairment may be due to confounding factors outside the explored pathways.

## Discussion

### Metabolic effects of the drug cocktail

It has previously been reported that female and male C57BL/6 mice treated with a drug cocktail of rapamycin, acarbose, and phenylbutyrate had an associated loss of weight with a matching trend in body fat mass (Jiang 2022). In longitudinal mouse studies, weight gain in male and female mice has been reported to increase as late as 22 months of age (Yanai 2021). AD has been associated with weight loss and the presence of Aβ-42 and its correlation with neuroinflammation has been a proposed mechanism for the inflammation’s effect on appetite (Sergi 2013, Sieske 2019). Additionally, frailty assessment in aging C57BL/6 mice has shown regardless of sex, frail mice had higher body weights and higher percentages of body fat which correlated with decreased endurance, slower walking speed, and less running wheel activity. Frail mice were also found to have decreased average lifespans (Baumann 2019). For female and male medicated cohorts, significant weight loss was observed in comparison to non-medicated cohorts which had non-significant weight loss over the study. It was also observed AAV-SHAM cohorts consumed more food on average compared to AAV-AD mice, indicating the drug cocktail may help to maintain body weight relative to the AAV-SHAM cohort. In addition, medicated cohorts had less fat mass prior to injection of the vector and at endpoint, apart from the male non-medicated AAV-SHAM cohort. Together, these trends suggest the drug cocktail improves body composition for both male and female mice, but preferentially in female mice.

Jiang also reported no significant difference in the consumption of the medicated feed (2022). Figure 3E-3F demonstrates a significant difference in consumption of the medicated feed regardless of vector injection. One of the hallmarks of aging is insulin resistance and poor glucose response. Acarbose prevents glucose absorption in the small intestine and rapamycin inhibits the mTOR pathway activated by glucose and their effects have been compared to caloric restriction (Harrison 2014, Blagosklonny 2019). Rapamycin has additionally been reported to increase energy expenditure (Makki 2014). Due to the activity of these drugs, the difference seen in consumption makes sense for medicated cohorts in that greater consumption would be seen to introduce more nutrients and reclaim homeostasis (Augustine 2020). Differences between consumption may also occur from a longer period of consuming the diet and the increased age of the cohorts.

Overall, any changes in body weight and fat mass align with findings in previous studies but prolonged exposure to the drug cocktail for mice past 22 months of age has not yet been characterized to the author’s knowledge. Additionally, this study demonstrates a robust impact of treatment with the drug cocktail on female cohorts and more subtle effect on male cohorts.

### Cognitive Impairment

The box maze assay has been validated for its ability to assess learning impairment in aging mice (Mukherjee 2019, Darvas 2020). It was previously reported that the drug cocktail reduced learning impairment in comparison to control cohorts (Jiang 2022). Female cohorts on average performed better in the box maze in comparison to male cohorts. In the separation of fast and slow learners for each cohort, it can be seen medicated slow learners on average performed faster than non-medicated slow learners. In addition, medicated fast learners performed faster than non-medicated fast learners with the same vector (Figure 3A-3C). In male cohorts, AAV-SHAM fast learners performed faster than AAV-AD fast learners. Surprisingly, the non-medicated AAV-SHAM cohort performed the fastest comparing their fast and slow learners against the other cohorts (Figure 3B-3D). In addition, the separation of fast and slow learners reveals a much more robust difference in performance for female cohorts than it does for male cohorts.

This is surprising as Jiang also reported faster escape times for male cohorts in comparison to female cohorts (2022) and it has been reported female mice specifically encounter spatial cognitive impairment earlier than males (Benice 2006). Treatment with rapamycin and acarbose has been shown to be effective if administered in mid- or late-life in mice (Herrera 2023). It was seen when normalizing food intake to body weight, male cohorts in comparison to their respective female cohorts, consumed significantly less food in the final four weeks before endpoint regardless of diet. In comparison of non-medicated male and female cohorts, food consumption normalized to body weight should be equivalent. The cognitive impairment phenotype may then be due to factors outside the primary anatomic region of analysis selected in this study (hippocampus). Regardless, the model still reflects the impact of the cocktail on cognitive impairment and suggests a vector dependent effect on performance.

### Expression of Aβ-42 and Tau

Alzheimer’s disease is a global disease of the human brain. However, the hippocampus is one of the first regions of the brain to show histopathological evidence of AD. Early-stage AD is characterized by mild cognitive impairment and the accumulation and aggregation of Aβ-42 and tau protein (Braak 1991). For this reason, the hippocampus was chosen to analyze the model. Green Fluorescent Protein (GFP) was included in the Aβ-42 construct as a reporter gene to indirectly validate the induction of the viral vector. For both male and female AAV-AD cohorts, density of positive expression of GFP was specific and significant in comparison to AAV-SHAM cohorts. Moreover, no differences were observed between the density of GFP in the hippocampus between the medicated and non-medicated cohorts. This comparison acted as a control for the potential effects of the drug cocktail on the vector.

In determining the expression of the human Aβ-42 sequence induced by the vector, the H31L21 antibody was chosen for its specificity. The commonly used antibody to detect Aβ pathology in human brains, 6E10, binds generally to amyloid proteins and demonstrated artifact cross reactivity when used in mouse brains. Female AAV-AD cohorts had significant in expression of Aβ-42 over AAV-SHAM cohorts regardless of diet, and the female medicated AAV-AD had a significant decrease in density compared to the non-medicated AAV-AD cohort. While AAV-AD specific density was seen in male cohorts, there was not a significant difference between the medicated and non-medicated AAV-AD cohort. This difference not only demonstrates the ability for the vector to enhance resilience to AD Aβ-42 accumulation, but also mirrors the reduction in cognitive impairment seen in the box maze. Common markers for phosphorylated tau such as AT8 and serine-404 target conserved sequences in mice. HT7 targets a sequence specific for human tau regardless of phosphorylation and so HT7 was chosen for human tau detection. Due to the only human tau sequence in the AAV-AD cohorts coming from the construct, specific staining was able to be achieved. Female AAV-AD cohorts saw significant and specific density of tau in comparison to AAV-SHAM cohorts. While on average male AAV-AD cohorts had specific density of tau, the differences between the AAV-SHAM cohorts were not significant. Additionally, male AAV-AD cohorts had half the tau density of AAV-AD female cohorts. This could be due to several AAV-AD males containing densities equivalent to their AAV-SHAM counterparts. Considering the viral vector used was the same for Aβ-42 and tau, and that male mice do seem to have higher levels of autophagy in comparison to female mice, it is possible an autophagic specificity for tau is present in male cohorts. It has previously reported tau is degraded via autophagy while current research on Aβ-42 clearance focuses on extra-cellular aggregates (Hamano 2021, Wei 2019, Bharadwaj 2020).

Overall, neuronal infection with the AAV-AD vector mediated specific and significantly higher density of protein expression of Aβ-42 and tau in the hippocampus of mice compared to the AAV-SHAM cohorts. Expression of Aβ-42 and tau demonstrates expression within neurons (Figure 4) indicating an early-stage model of AD, as later stages are characterized by extracellular Aβ-42 aggregates, not seen during pathologist review of H&E-LFB slides.

### Neuropathology and Non-Neuronal Cells

The progression of AD even in the early stages affects multiple processes resulting in a more severe “aging” phenotype. One of the first affected pathways tied closely with the onset of mild cognitive impairment is synaptic integrity. It has been suggested that soluble Aβ-42 causes synaptic dysfunction prior to the formation of plaques and clinical detection (Solkoe 2002, John 2022). In female medicated cohorts, increased levels of synaptophysin were seen when compared to non-medicated cohorts. In addition, increased levels of PSD-95 were also seen in medicated cohorts when compared to non-medicated cohorts of the same vector. In comparison with the AAV-AD and AAV-SHAM cohorts, synaptophysin and PSD-95 were decreased in AAV-AD cohorts relative to the respective AAV-SHAM cohorts. These results suggest the drug cocktail improves synaptic integrity, while the presence of AAV-AD proteins harm synaptic integrity. These findings are mirrored in the box maze data where the female medicated AAV-SHAM cohort showed less learning impairment. Male cohorts saw similar trends, with a smaller difference in synaptophysin between AAV-AD cohorts and greater expression of PSD-95 in non-medicated cohorts. The drug cocktail appears to enhance synaptophysin expression in medicated mice but not the expression in PSD-95. It has been previously found that PBA reduces cognitive impairment and increased synaptic plasticity in mouse models of AD (Ricobaraza 2009, Wiley 2011). Male non-medicated cohorts seem to have higher densities of PSD-95 and synaptophysin in comparison to female non-medicated cohorts. Given the greater burden of Aβ-42 in non-medicated female mice, the findings of the study may suggest these changes are driven by AD proteins, though the differences in box maze suggest synaptic integrity across sexes is dependent on more than synaptophysin and PSD95.

The buildup of Aβ-42 and tau protein aggregates can cause neuronal injury through accumulated double strand breaks (Shanbhag 2019) and a vicious cycle of pro-inflammatory cytokines (Sinyor 2020). Additionally, these increases activate an autophagic response which has been reported to clear Aβ-42 in mouse models of AD (Heckmann 2019). For both male and female cohorts, medicated cohorts had less inflammation in comparison to non-medicated cohorts with the same vector, however, the non-medicated AAV-AD cohort had a much more substantial level of inflammation. Overall, it does appear the male cohorts have higher levels of inflammation in comparison to the female cohorts. The drug cocktail has previously been validated for its ability to reduce inflammation in major systemic organs as well as in the brain (Jiang 2022, 2023) matching our study results. The level of inflammation suggests an AAV-AD specific instigation of inflammation and the drug cocktail’s ability to modulate that response. Female AAV-AD cohorts had more DNA damage in comparison to their AAV-SHAM counterparts and medicated cohorts had less DNA damage in comparison to non-medicated cohorts. The male medicated AAV-AD cohort has less DNA damage than the non-medicated cohorts, however the medicated AAV-SHAM cohort had more DNA damage than the non-medicated AAV-SHAM cohort. The male cohort again has a lower burden of Aβ-42 in the brain and lower protein aggregation in the neurons may reduce neuronal injury in the medicated cohort. Additionally, the non-medicated AAV-SHAM performed well in the box maze suggesting less progressive DNA damage may contribute to its faster learning times. In female cohorts, autophagy was increased in for the medicated AAV-SHAM cohort in comparison to the non-medicated AAV-SHAM cohort, but this trend was inverted for AAV-AD cohorts. In males, non-medicated cohorts had higher levels of autophagy in comparison to medicated cohorts. Female cohorts did see a significant decrease in Aβ-42 expression which may explain the decrease in autophagy for the medicated AAV-AD cohort. Males see the inversion of what is expected, though autophagy does mirror trends in PSD95 which may indicate opposing sex differences in hippocampal health.

Microglia and astrocytes are controversial for their role in AD pathogenesis. While microglia help to degrade Aβ-42, they also participate in the release pro-inflammatory cytokines which activate astrocytes, preventing them from supporting synapse plasticity and maintenance (Liddelow 2017) as well as participating in synapse loss through engulfment (Hansen 2018). Astrocytes participate in synapse maintenance, regulation of the inflammatory response and provide neurons with nutrients but hyperproduce hydrogen peroxide and atrophy as AD progresses (Chun 2020). In our study, female medicated cohorts had increased levels of microglia in comparison to non-medicated cohorts. Male cohorts had the opposite trend where medicated cohorts had less activated microglia expression in comparison to non-medicated cohorts with the same vector. For male cohorts, the trends in microglia mirror autophagy, suggesting the levels of autophagy seen may be activated by microglia. The female cohorts demonstrated the opposite trend of what was expected. It is possible microglia, in combination with a reduction in inflammation, lead to more effective response to increases in Aβ-42, which would explain increases in microglia and trends in Aβ-42 expression for male and female cohorts. In female cohorts, astrocytes have increased density in medicated cohorts when compared against non-medicated cohorts. Male mice show a mirrored trend within cohorts of the vector. Activation of microglia in females could be the reason for the activation of astrocytes as the trends are mirrored across cohorts. As previously stated, sex differences in the activation response of astrocytes may explain the differences between non-medicated cohorts.

Overall, the drug cocktail appeared to rescue AAV-AD inflicted degeneration of synaptic integrity, increased inflammation and DNA damage, and enhance healthy microglia-neuron interactions.

### Associations

For non-medicated females, autophagy activated by microglia was most likely responsible for faster box maze times resulting in the negative associations. Aβ-42 and tau associating strongly implies consistent relative expression of both proteins and the association of both with inflammation implies specific responses to the AAV-AD vector. Postsynaptic integrity had negative associations with inflammation and autophagy likely as a response to production of inflammatory proteins and the necessity to degrade excess proteins. These associations demonstrate the wide necessity of autophagy and the far-reaching impact of inflammation in response to the AD protein expression.

In contrast, female medicated cohorts demonstrate positive associations with Aβ-42 and DNA damage are the driving factors for box maze, with DNA damage associating with inflammation. The overall reduction in inflammation is likely from a reduction in expression of Aβ-42, with inflammation arising from DNA damage instead. Mice with less Aβ-42 and DNA damage then perform better in the box maze. Aβ-42 and tau have consistent relative expression as well as a strong association with activated astrocytes. The association with activated astrocytes may be a compensatory measure against an increase in tau as it appears tau and microglia have a strong negative relationship. These associations suggest that Aβ-42 and tau play a role in the activation of astrocytes.

Between the non-medicated and medicated females, differences induced by the medicated diet are revealed. Non-medicated female cohorts have much greater amounts of inflammation and upregulation of autophagy induced by Aβ-42 and tau. In contrast, medicated mice over all do not have the same up-regulation of inflammation and autophagy but do display an increase in synaptic integrity as well as active astrocyte association with Aβ-42 and tau expression. The drug cocktail thus demonstrates mediation of synaptic integrity, inflammation, and autophagy within the female model.

Box maze in male non-medicated cohorts had strong negative associations with presynaptic integrity and inflammation. Tau had strong associations with Aβ-42, implying consistent relative expression, and inflammation. Inflammation and autophagy induced by microglia had a strong association as a response to AAV-AD protein expression. Lastly, microglia had a strong negative association with presynaptic integrity, possible due to engulfment of synapses. These associations suggest an inflammatory cascade response to the AAV-AD vector ending in cognitive impairment.

Medicated males had a strong negative association with Aβ-42 and box maze. This is a counter intuitive relationship. Microglia have a strong association with Aβ-42. There does not seem to be astrocyte activation and so microglia carry the burden for the residual inflammation caused by Aβ-42. Tau had a strong association with DNA damage, and presynaptic integrity had a strong negative association with inflammation. These associations imply Aβ-42 induces a microglia response. Due to the similarity in the box maze, it appears in the medicated cohorts there is no effect of the AAV-AD on box maze performance.

The differences between the males demonstrate small effects of the drug cocktail. Similar to the female cohorts, the drug cocktail does reduce inflammation, autophagy, and improves synaptic integrity. However, it does not appear like in the female cohorts, improvement of synaptic integrity results in a reduction of cognitive impairment. This could be due to additional variables outside the hippocampus impacting overall performance.

In an evaluation of the model, resilience to aging was enhanced in male and female mice primarily in pathways of autophagy, synaptic integrity, and inflammation, with a much bigger increase to resilience seen in female mice. While cognitive impairment appeared to be subtle, Alzheimer’s disease is a disease of slow progression where clinical phenotypes present after neuropathology has been well established. Additionally, the drug cocktail significantly delayed the onset the presence of Aβ-42 in female mice. It is important to note the effects seen in this study were analyzed as a combination of all three drugs and individual drugs were not tested in this study. Jiang et al. demonstrated a stronger effect in increasing resilience in mouse brains for the combination of drugs (2023). More in depth analysis of specific downstream targets and pathways of each drug included in the cocktail would need to be evaluated for determining their contribution to the results. In addition, while young controls have been used in the past to validate the effects of the drug cocktail (Jiang 2023), a control cohort of mice not infected with either vector may help to elucidate a baseline of aging pathology and injection effect within the AAV cohorts. All cohorts in this study were infected with an AAV vector, so any plausible effect of the virus would be controlled for. This model of early-stage Alzheimer’s disease is a valuable research tool not only for exploratory research but also for evaluation of AD expression within the context of aging.

In conclusion, female mice were more susceptible to the early development of AD neuropathology and learning impairment, and more responsive to treatment with the drug cocktail in comparison to male mice. Translationally, a model of AD where females are more susceptible would have greater value as women have a greater burden and incidence of disease compared to men. Future studies will look to establish genetic tests for biological aging and similar in-depth analysis of neuropathology in additional regions of the brain.

## Acknowledgments

This work was supported by National Institutes of Health grants R01 AG057381 (Ladiges and Darvas, coPI’s) and R01 AG057381 (Ladiges PI).

The California Institute of Technology (Caltech) has generously allowed the AAV vectors, based on AAV-PHP and/or variants, AAV-PHP.B, AAV-PHP.B2, AAV-PHP.B3, AAV-PHP.eB, AAV-PHP.S, to be distributed with the understanding that requesting investigators need to acknowledge Caltech in any publications in which AAV-PHP was used, specifically Viviana Gradinaru, Ph.D. and Benjamin E. Deverman, Ph.D., and cite Chan et al., 2017 Nat Neurosci, 20(8):1172-1179. (Pubmed) for AAV-PHP.eB and AAV-PHP.S and/or Deverman et al., 2016 Nature Biotechnology 34: 204-209 for vectors AAV-PHP.B, AAV-PHP.B2 and AAV-PHP.B3. AAV-PHP is a modification of AAV9 provided by the University of Pennsylvania to Caltech.

